# FOXM1 acts sexually dimorphically to regulate functional β-cell mass

**DOI:** 10.1101/2023.01.12.523673

**Authors:** Guihong Peng, Elham Mosleh, Andrew Yuhas, Kay Katada, Christopher Cherry, Maria L. Golson

**Author notes:** Corresponding Author: Maria L Golson.

## Abstract

The transcription factor FOXM1 regulates β-cell proliferation and insulin secretion. Our previous work demonstrates that expressing an activated form of FOXM1 (FOXM1*) in β cells increases β-cell proliferation and mass in aged male mice. Additionally, FOXM1* enhances β-cell function even in young mice, in which no β-cell mass elevation occurs. Here, we demonstrate that FOXM1 acts in a sexually dimorphic manner in the β cell. Expression of FOXM1* in female mouse β cells does not affect β-cell proliferation or glucose tolerance. Transduction of male but not female human islets with FOXM1* enhances insulin secretion in response to elevated glucose. Estrogen contributes to diabetes susceptibility differences between males and females, and the estrogen receptor (ER)α is the primary mediator of β-cell estrogen signaling. We show that FOXM1* can rescue impaired glucose tolerance in female mice with a pancreas-wide ERα deletion. Further, FOXM1 and ERα binding sites overlap with each other and with other β-cell-enriched transcription factors, including ISL1, PAX6, MAF, and GATA. These data indicate that FOMX1 and ERα cooperate to regulate β-cell function and suggest a general mechanism contributing to the lower incidence of diabetes observed in women.

## Introduction

Despite an elevated obesity prevalence in women, men exhibit a higher prevalence of diabetes until around age 65 (1,2). The same contrast between male and female humans also occurs in mouse models; usually, female mice are resistant to glucose intolerance and diabetes in models that result in diabetes in males (3,4), with a few exceptions such as the β-cell-specific *Hnf4α* knockout and global *Prlr* knockout (5-7).

This resistance to developing impaired glucose homeostasis likely arises from estrogen action in peripheral tissues and in female β cells. Supporting this conclusion, oophorectomized female mice develop glucose intolerance and diabetes at similar levels as male mice (8). In addition, female mice lacking the estrogen receptor α (ERα) specifically in the pancreas have impaired glucose tolerance (9) and their β cells are more susceptible to apoptosis in response to oxidative stress (10). Moreover, *in-vitro* 17β-estradiol treatment of islets results in increased insulin transcription, content, and release (11), while knockdown of ERα reduces insulin secretion (10). Finally, ERα^-/-^ mice display compromised embryonic β-cell proliferation and blunted β-cell mass recovery after partial pancreatectomy (12).

The forkhead box transcription factor FOXM1 promotes cell division and insulin secretion in the β cell. At least in young mice, FOXM1 is upregulated in response to stimuli that promote β-cell replication, including pregnancy, partial pancreatectomy, and obesity (4,13,14). Both male and female mice that lack FOXM1 in the pancreas (*Foxm1*^*Δpanc*^ mice) exhibit a ∼60% diminished β-cell mass at nine weeks (4,15). This diminished β-cell mass translates to glucose intolerance or frank diabetes in *Foxm1*^*Δpanc*^ males. However, *Foxm1*^*Δpanc*^ female mice maintain normal glucose homeostasis unless subjected to additional metabolic stress, such as pregnancy (4,13).

*Foxm1* expression diminishes with age, concomitant with decreased basal and stimulated β-cell proliferation (16). Using a model in which an activated form of FOXM1 is expressed in β cells (*RIP-rtTA*;*Tet-O-HA-Foxm1*^*ΔNRD*^ mice, referred to hereafter as β-FoxM1* mice), we previously showed that replenishing *Foxm1* in aged male β cells increases β-cell mass and improves glucose tolerance (16). In addition, male β-FoxM1* β cells are resistant to streptozotocin- and cytokine-induced β-cell death (17).

At two months of age, when no increase in β-cell mass was observed, β-FoxM1* males also demonstrated improved glucose tolerance (16). β cells expressing FoxM1* displayed increased Ca^2+^ entry in response to elevated glucose. Moreover, examination of size-matched islets from male mice lacking FOXM1 specifically in β cells revealed diminished insulin secretion per β cell in the absence of FOXM1.

In the current study, single-cell RNA sequencing (scRNAseq) in male β-FoxM1* mice demonstrated an elevated expression of genes and pathways involved in β-cell proliferation, insulin secretion, and apoptosis. We expressed FOXM1* in human islets and observed increased insulin secretion in male but not female islets, prompting an investigation that revealed a sexually dimorphic action for FOXM1 in β cells. Because FOXM1 and ERα regulate similar β-cell functions—proliferation, insulin secretion, and apoptosis—and because FOXM1 and ERα physically cooperate in other cell types (18), we investigated the interaction of FOXM1 and ERα in β cells.

## Research Design and Methods

### Mice

*RIP-rtTA* (19), *Tet-O-HA-Foxm1*^*ΔNRD*^ *(17), Tet-O-Cre* (20), and *ERα*^*fl*^ (21) mice and genotyping have been described previously. Mice were maintained on a C57Bl6/J background and housed either at the University of Pennsylvania or at Johns Hopkins University with a 12-hour light-dark cycle. At eight weeks, mice were treated with 2% doxycycline (Sigma-Aldrich, St. Louis, MO) in drinking water for two weeks before experimental procedures. Littermate controls were included in all experiments. All procedures were conducted following the Institutional Animal Care and Use Committee guidelines at the University of Pennsylvania or Johns Hopkins University.

### Glucose tolerance tests

Glucose tolerance tests (GTTs) were performed as previously described (22).

### Mouse islet isolation

Mouse islets were isolated by collagenase digestion, followed by Ficoll gradient and handpicking.

### Perifusions

Perifusions were performed by the Islet Cell Biology Core and insulin measured by the Radioimmunoassays and Biomarkers Core at the University of Pennsylvania.

### FoxM1* adenovirus generation and transduction

The *Foxm1* adenovirus plasmid was generated as previously described (23). In brief, a mouse *Foxm1* cDNA lacking the self-suppressing N-terminal domain (17) was inserted into the multi-cloning site of an adenoviral vector downstream of a TRE element. This insertion site is upstream of cDNA encoding enhanced green fluorescence protein (*eGFP*) and reverse tetracycline transactivator (*rtTA*) separated by 2A peptide under the control of the rat *insulin* promoter *(RIP)*. Vector Biolabs (Malvern, PA) grew and expanded the encoded adenovirus. The eGFP adenovirus was purchased from Vector Biolabs. Human islets obtained from the Integrated Islet Distribution Program (IRB waiver IRB00239926) were treated with 0.05% trypsin for 3 minutes at 37°C, then transduced using spinfection with 100 islets per microcentrifuge tube at a multiplicity of infection (MOI) of 200 for 45 minutes at 800 rpm. Islets were then cultured in Prodo (Aliso Viejo, CA) media supplemented with 5% human serum and 1 µg/mL doxycycline (Sigma-Aldrich) for three days before perifusions.

### Immunofluorescence, immunohistochemistry, and quantification of histological features

Pancreata were fixed and embedded in paraffin, and immunofluorescence quantification was performed as previously described (22). For β-cell mass, at least five sections taken throughout the pancreas were immunolabeled against Insulin using DAB visualization (Jackson ImmunoResearch Laboratories, West Grove, PA). DAB and Eosin areas were quantified with QuPath (24). β-cell mass was calculated as previously described (25). Antibodies used included guinea pig anti-insulin (1:50-1:500, Dako, now Agilent) and mouse anti-Ki67 (1:500). Secondary antibodies were used at 1:400-1:600 and obtained from Jackson ImmunoResearch Laboratories.

### Chromatin immunoprecipitation sequencing

βTC6 cells were cultured in 5 mM DMEM supplemented with 10% hormone-depleted fetal bovine serum for one week before treating with vehicle or 100µM 17β-estradiol (Sigma-Aldrich) in ethanol (final concentration 0.1%) for one hour. Chromatin was collected as previously described (26) and sonicated (Covaris) to recover fragments between 200-800 bps. ChIP was performed on five replicates per treatment group, as previously described (26). Antibodies included rabbit anti-FOXM1 (Diagenode, Denville, NJ), goat anti-FoxA2 (Santa Cruz, Dallas, TX), rabbit anti-ERα (Santa Cruz), and a rabbit IgG control (Santa Cruz). The Hopkins University Single-Cell and Transcriptomics Core performed library prep and sequencing. Quality control and adapter trimming of fastq files were performed using fastp (27). Bowtie2 was used to align fastq files against the GRCm39 reference genome (28). Genrich was used for consensus peak calling. Differential accessibility analysis was performed using DESeq2 after merging peaks across treatment groups and determining counts within each peak by sample (29). ChIPpeakAnno was used to annotate differentially accessible peaks. HOMER was used to identify binding motifs (30).

### Single-cell RNA sequencing

The University of Pennsylvania Functional Genomics Core performed library prep and RNAseq as previously described (31). Alignment and preliminary quality control were performed using Cellranger v6.0.1 from 10X Genomics (Pleasanton, CA) with a customized reference genome, including transgenes, built from the included GRCm38 references. The resulting count files were filtered using a UMI threshold of 500 transcripts, a feature threshold of 100 genes, and a mitochondrial transcript threshold of 25% contribution from mitochondrial RNA. Normalization, scaling, principal component analysis, uniform manifold approximation and project (UMAP), and clustering were performed using Seurat (32). Batch effect correction of principal components was performed using Harmony (33). Clusters were identified using a combination of differential expression comparing each cluster to all other cells and marker gene expression. Subsequent differential expression was also analyzed by comparing treatment groups within each cluster. Mann-Whitney U tests were performed for all differential expression with Benjamini Hochberg procedure for multiple testing correction. Gene set analysis of all upregulated genes listed in Table 1 was conducted using DAVID (34).

**Table 1.**
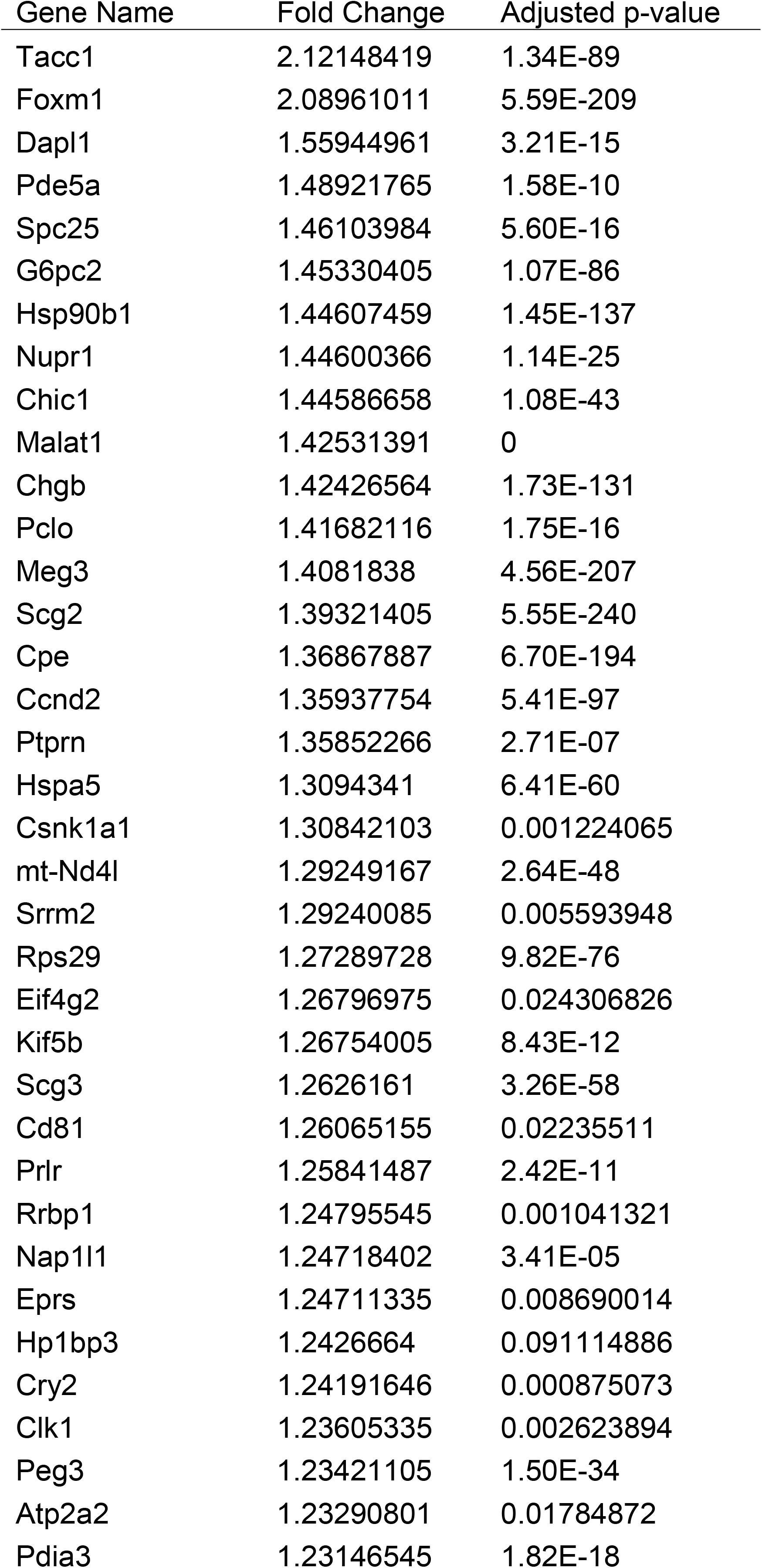
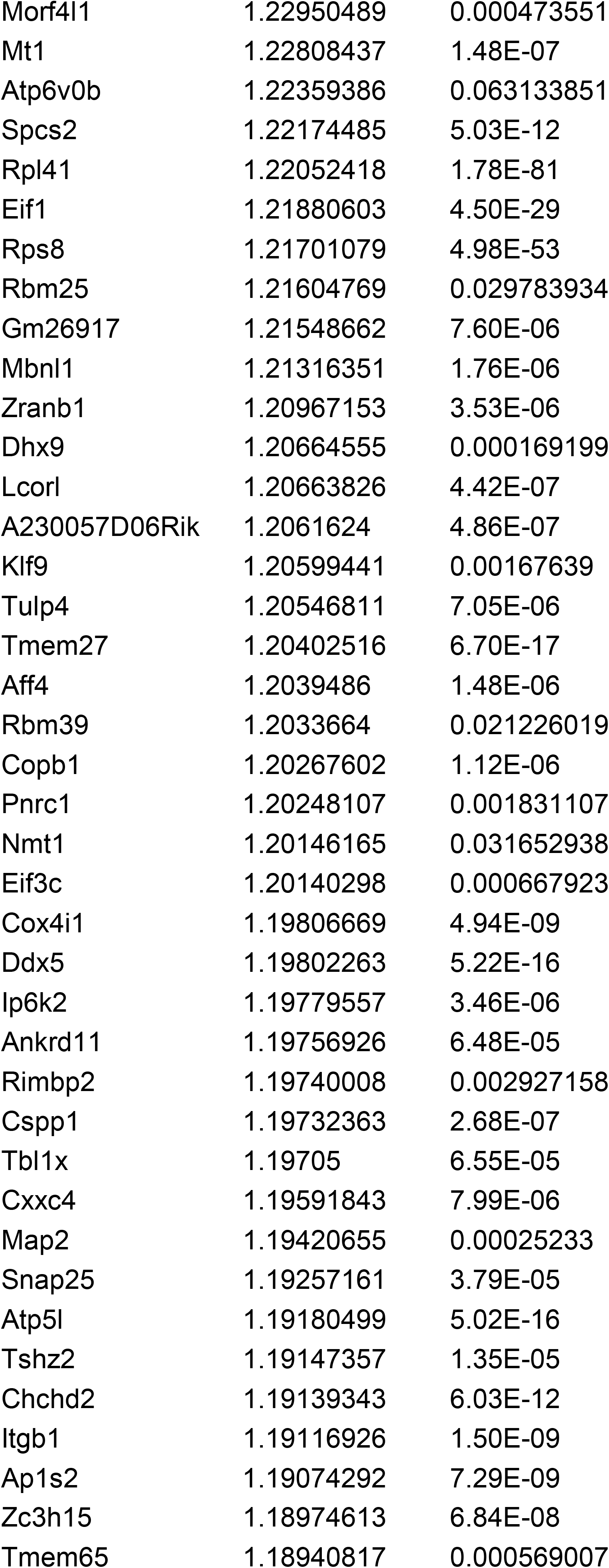
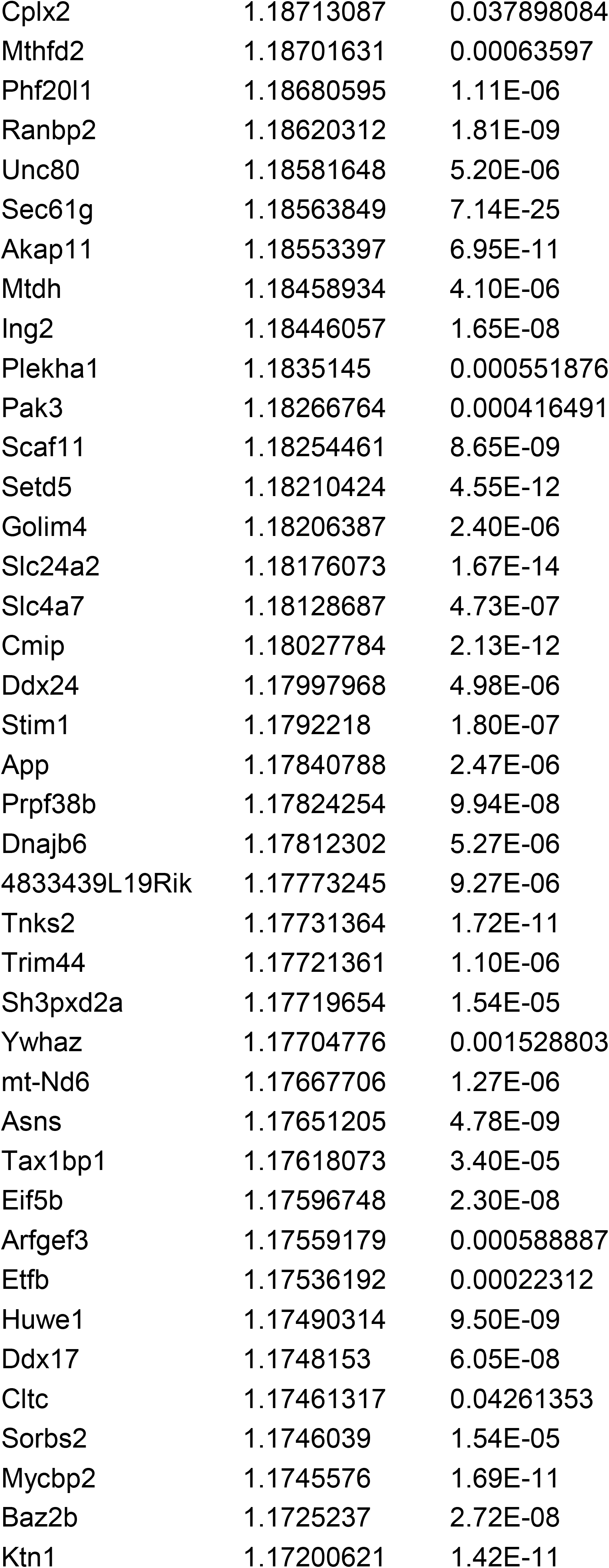
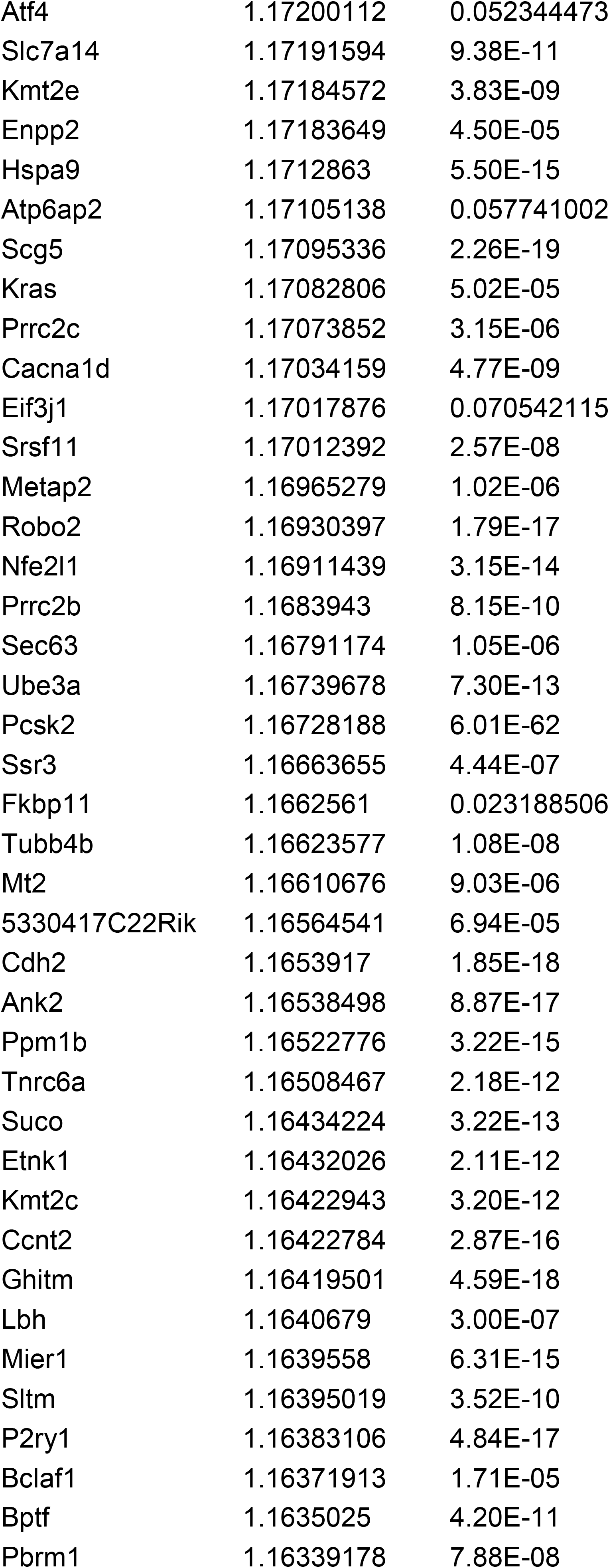
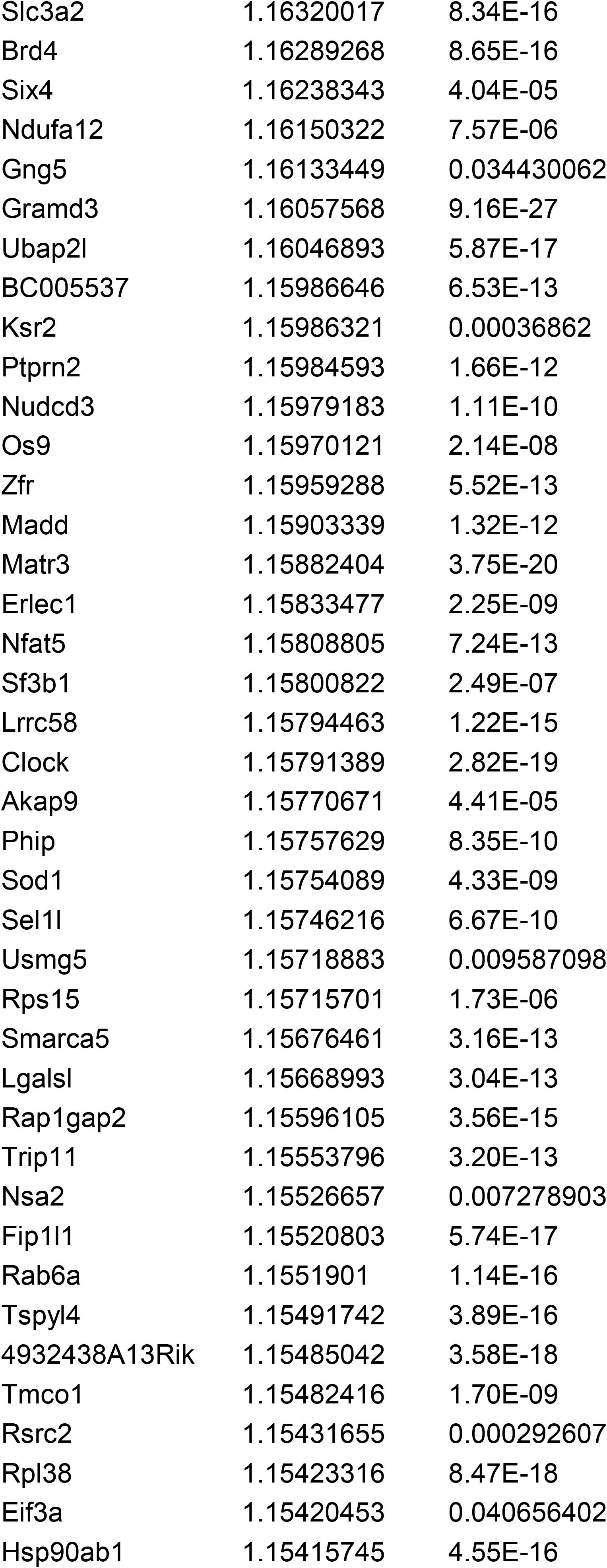
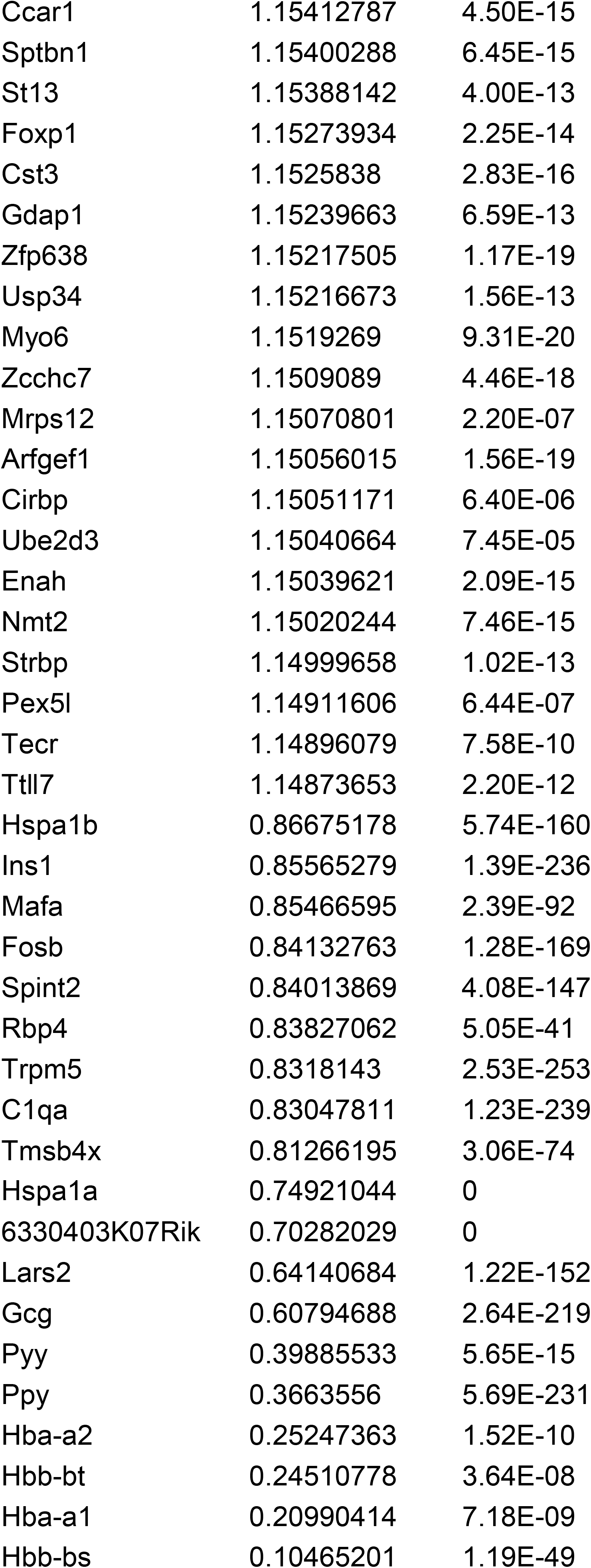
Differentially regulated genes in male β-FoxM1 β cells.

### Statistical analysis

For experiments other than RNAseq and ChIPseq, significance was analyzed using ANOVA, Two-Way ANOVA, or multiple t-test analysis, as appropriate, using Prism software (GraphPad, San Diego, CA). Error bars are SD.

## Results

### scRNAsequencing reveals increased expression of genes involved in insulin secretion and β-cell function

We previously performed RNA sequencing on whole islets from male β-FoxM1* mice and reported downregulation of pro-inflammatory cytokines and interleukins (16). To refine these results to gene expression differences solely in β cells, we performed scRNAseq. Differentially regulated genes are listed in Table 1. The top 20 upregulated genes in male β-FoxM1* β cells primarily either promote proliferation—*Tacc1, Foxm1, Spc25, Ccnd2, Prlr*, and *Pde5a—*or regulate insulin processing or secretion—*G6pc2, Chrb, Pclo, Meg3, Scg2*, and *Scg3—*or protect against apoptosis—*Hsb90b1, Nupr*, and *TALAT* (Figure 1A). DAVID pathway analysis revealed the enrichment of gene sets involved in protein processing, protein export, oxidative phosphorylation, and insulin secretion, among others (Figure 1B).

**Figure 1.**
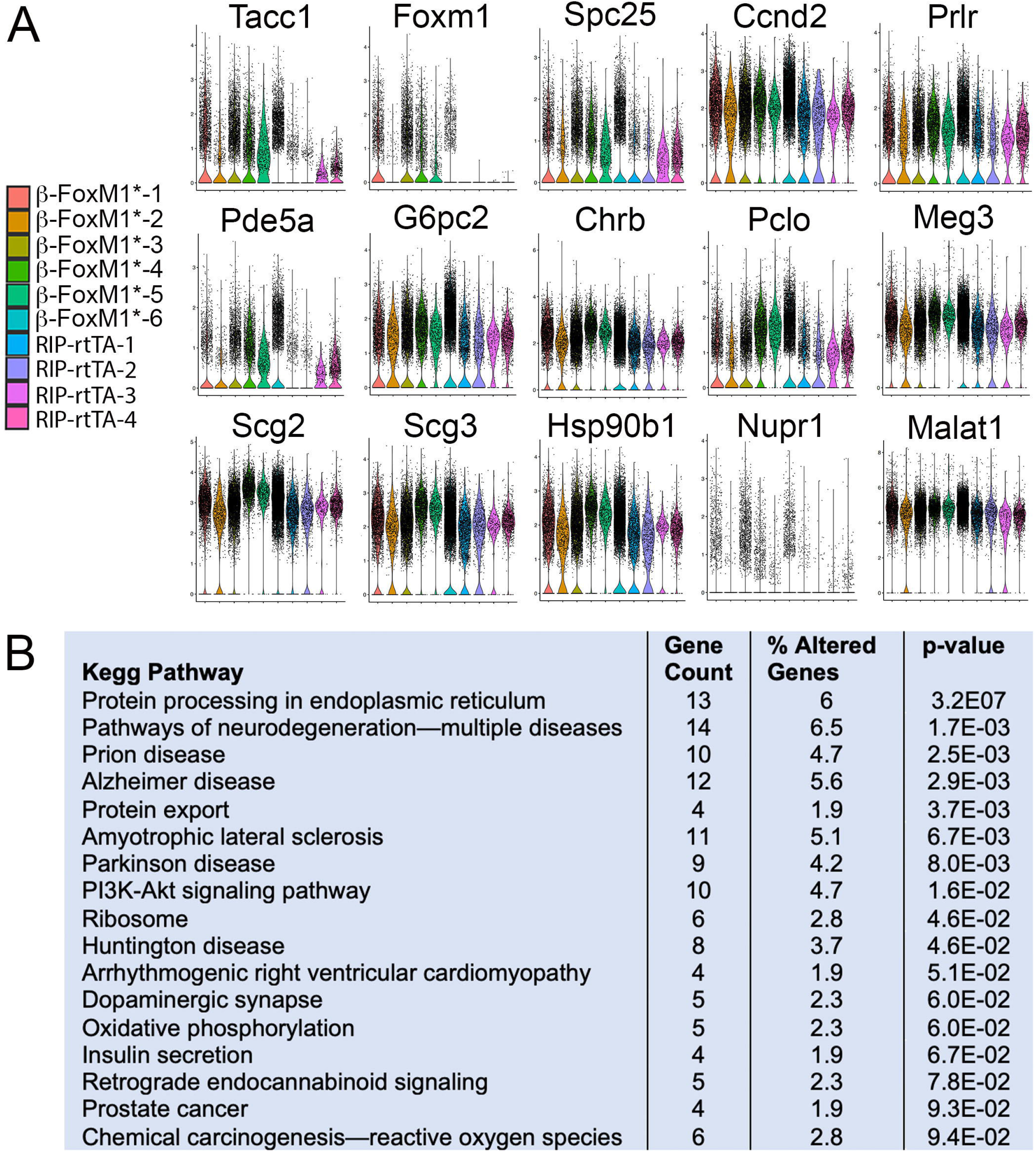
Single-cell RNAseq analysis of β cells in male β -FoxM1 * mice. A) Violin plots of genes of interest that are upregulated by >1.25-fold and have an adjusted p-value <1.58E-10. Genes displayed are in the top 20 upregulated genes and promote β-cell proliferation or β -cell function or protect against apoptosis. Within each functional group, described in text, genes are ordered by fold-change. B) DAVID analysis of KEGG pathways.

### Activated FOXM1 expression has different effects in male and female islets

Because we previously demonstrated that activated FOXM1 expression can improve male β-cell function, we next investigated whether FOXM1* could increase insulin secretion in human islets. Male human islets transduced with an adenovirus encoding *Foxm1*^*ΔNRD*^ driven by the RIP promoter secreted more Insulin in response to high glucose, but not to IBMX, than did islets transduced with GFP (Figure 2, A-B). Surprisingly, though, expressing FOXM1* in female human islets did not alter insulin secretion (Figure 2, C-D).

**Figure 2.**
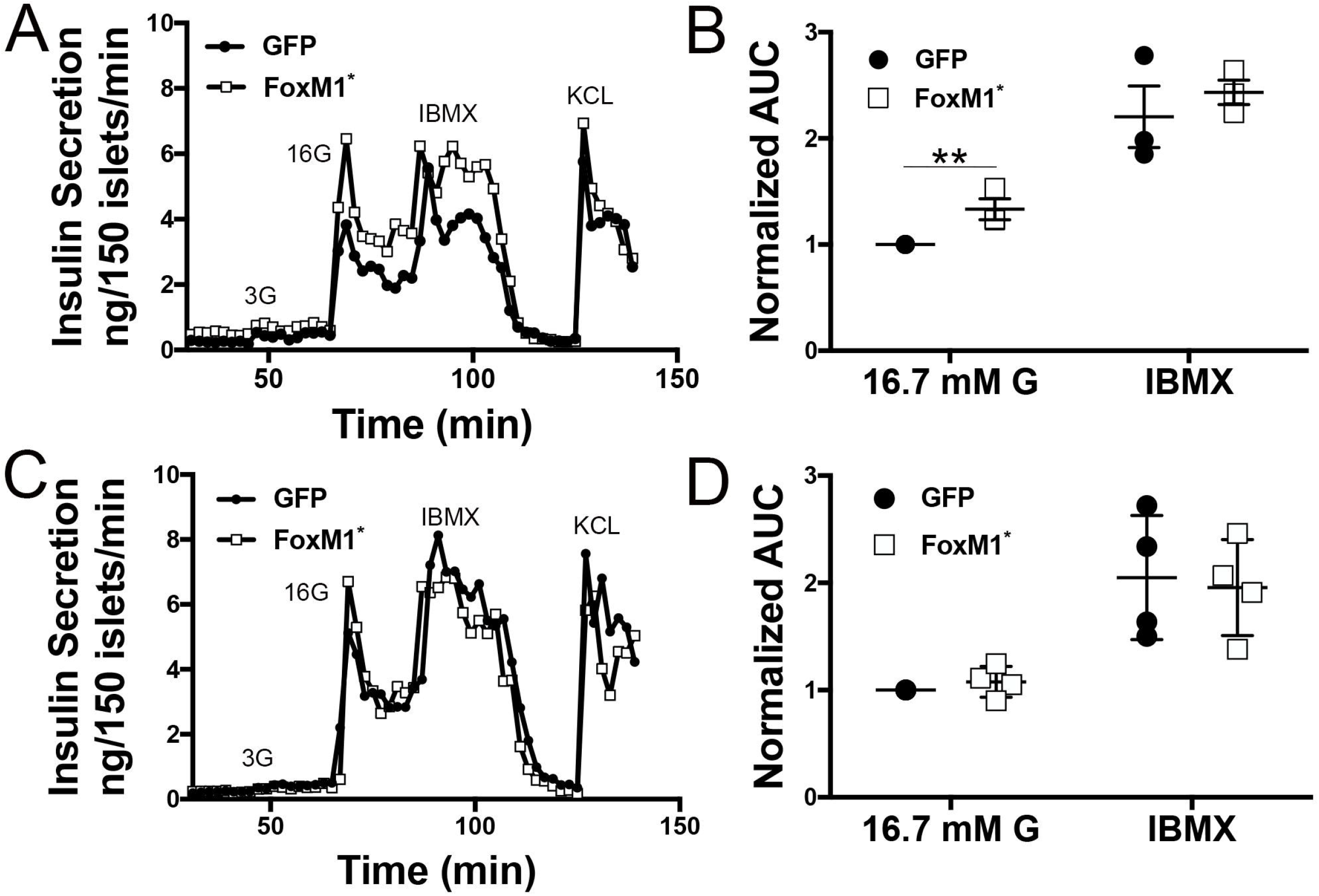
Human male but not female islets transduced by activated FoxM1 secrete more insulin in response to high glucose than control islets. A-D) Representative perifusions (A,C) and insulin secretion AUC in response to high glucose and IBMX, normalized to high glucose AUC (B,D) from male (A,B) and female islets (C,D).

Because of this discrepancy in insulin secretion potentiation by FOXM1* between male and female human islets, we explored whether there was sexual dimorphism between male and female β-FoxM1* mice. FOXM1 activation did not result in increased β-cell mass or proliferation compared to *RIP-rtTA* controls in either male (16) or female 10-week-old β-FoxM1* mice, even when females were challenged by pregnancy and examined at gestational day (GD)15.5, a time point at which β-cell proliferation is high and (Figure 3, A-J and). However, in contrast to the beneficial effects seen in male β-FoxM1* mice (16), activated FOXM1 did not increase β-cell mass or proliferation in aged female mice (Figure 3, K-P). In addition, unlike male β-FoxM1* mice (16), neither young (virgin or pregnant) nor aged β-FoxM1* female mice demonstrated improved glucose tolerance (Figure 3, Q-S).

**Figure 3.**
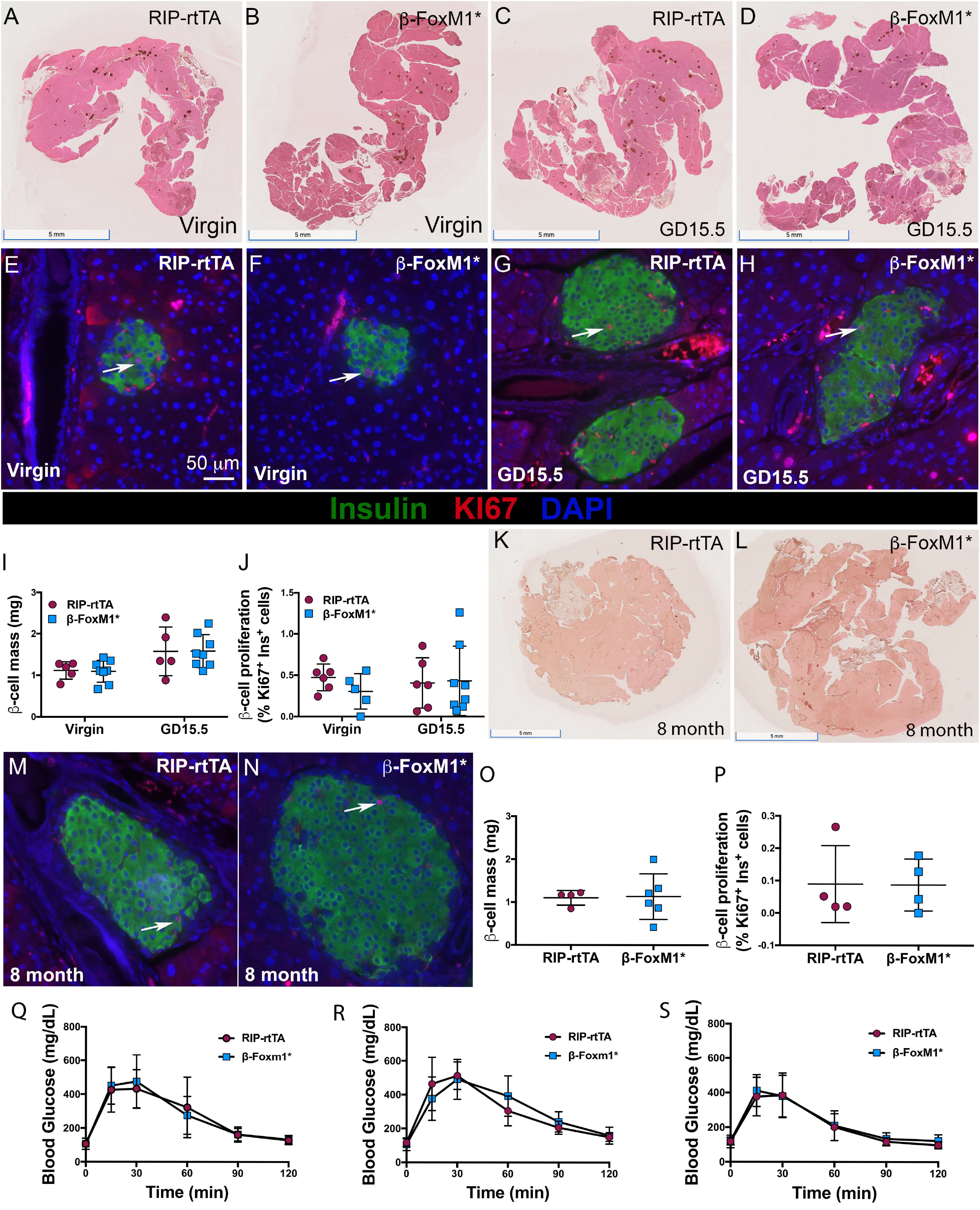
(β-cell mass and proliferation and glucose tolerance are unchanged in young and aged female ✉-FoxM1* mice. A-D) Representative images for insulin immunolabeling on two-month-old virgin (A-B) or GD15.5 (C-D) pancreata. E-H) Representative images for insulin and Ki67 immunofluorescence on two-month-old virgin (E,F) or GD15.5 (G,H) pancreata. (I) Quantification of [β-cell mass in two-month-old [β-FoxM1* female mice. (J) Quantification of [β-cell proliferation in two-month-old (β-FoxM1* female mice. K-L) Representative images for insulin immunolabeling on eight-month-old virgin pancreata. M-N) Representative images for insulin and Ki67 immunofluorescence on eight-month-old virgin pancreata. Pseudocolors used are the same as for (E-H). (O) Quantification of β-cell mass in eight-month-old β-FoxM1* female mice. (P) Quantification of [β-cell proliferation in eight-month-old p-FoxM1* female mice. Q-S) Glucose tolerance tests for (Q) two-month-old virgin β-FoxM1* mice, (R) two-month-old GD15.5 (β-FoxM1* mice, and (S) eight-month-old virgin p-FoxM1*.

### Activated FoxM1 rescues glucose tolerance defects in ERα^Δβ^ female mice

ERα is the predominant estrogen receptor expressed in β cells (35). We, therefore, postulated that removing ERα from female β cells would transform them to a more “masculinized” state, in regards to ERα target binding, and that activated FOXM1 could improve glucose in female mice ERα in β cells, just like it had in otherwise wild-type male mice (16). To test this hypothesis, we ablated *Esr1* in β cells (ERα^Δβ^ mice) using the *RIP-rtTA*;*Tet-O-Cre* system. ERα^Δβ^ females displayed mildly impaired fasted glucose and glucose tolerance, both of which were rescued by adding activated FOXM1 to ERα^Δβ^ β cells (β-FoxM1*;ERα^Δβ^ mice; Figure 4A-C). ERα^Δβ^ females also displayed increased β-cell mass, which again was corrected by the presence of FoxM1* (Figure 4D). Both ERα^Δβ^ mice and β-FoxM1*;ERα^Δβ^ mice displayed elevated body weight compared to control and β-FoxM1* mice, but no significant difference was apparent between ERα^Δβ^ and β-FoxM1*;ERα^Δβ^ females (Figure 4E).

**Figure 4.**
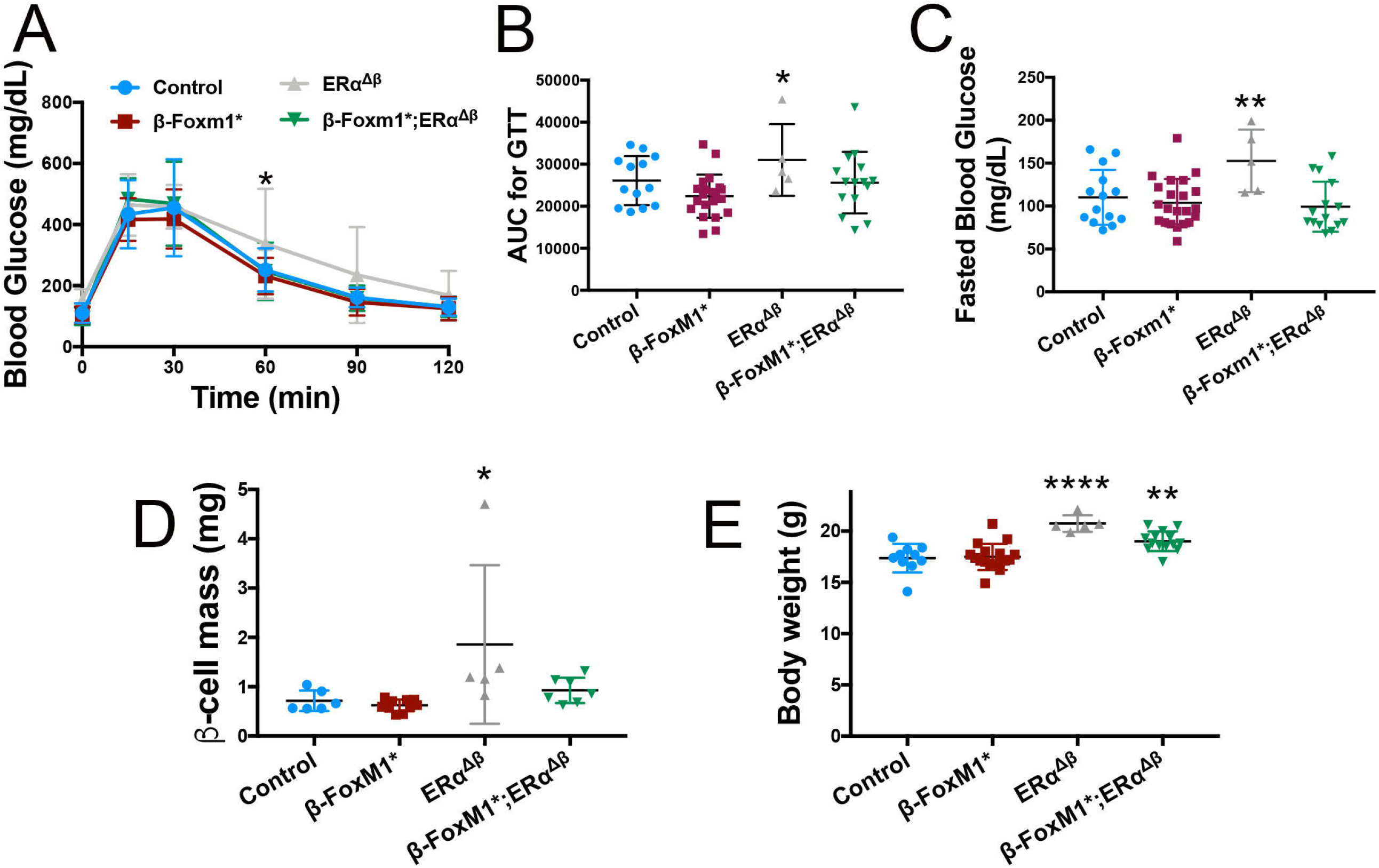
Activated FoxM1 rescues glucose tolerance defects in female ERα mice. A-C) Glucose excursion during a GTT (A-B) and fasted blood glucose (C) are elevated in female mice lacking ERα in β cells, but both glucose excursion and fasted glucose are normalized in ERα mice that also express activated FoxM1 in β cells. n=S-22 for both GTTs and fasted blood glucose. (C) β-cell mass is elevated in ERα but normalized by the presence of FoxM1. (D) Both ERα and β-FoxM1 *;ERα mice display elevated body weight. In (A-D), all mice have at least one allele of RIP-rtTA.

### FOXM1 and ERα binding sites overlap in a β-cell line

Since FOXM1, FOXA1, and ERα physically cooperate in breast cancer cells to drive the expression of multiple cell-cycle genes (18), we investigated whether FOXM1, ERα, and FOXA2 are also recruited to the same chromatin locations in β cells. We chose to examine FOXA2 instead of FOXA1, as FOXA2 is required for β-cell differentiation and identity maintenance, while FOXA1 has a limited role in β-cell development and is severely downregulated in mature β cells (36-38). ChIP was performed against FOXA2, FOXM1, and ERα on chromatin from βTC6 cells grown in media supplemented with hormone-depleted FBS to remove serum estradiol and treated with either 17β-estradiol (E_2_) or vehicle (Figure 5).

**Figure 5.**
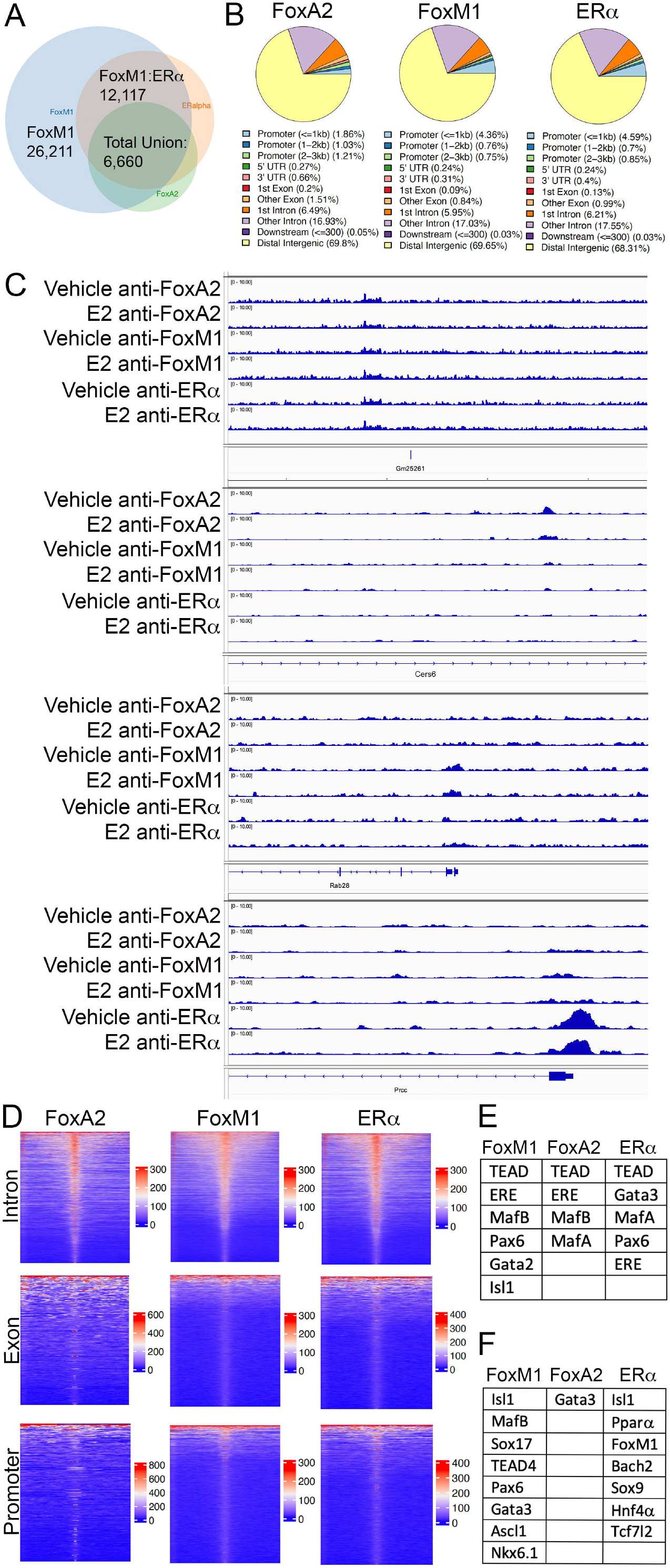
F0XM1, F0XA2, and ERα binding sites overlap in □ TC6 cells treated with 17-β-estradiol. A) Venn diagram of F0XM1, F0XA2, and ERα binding-site overlap B) Categorization of binding site locations of F0XM1, F0XA2, and ERα. C) Examples of sites bound by F0XA2, F0XM1, and ERα. Each track represents the average of 5 independent ChIP experiments. Sites shown include intergenic regions, promoters, and introns. D) Heatmap of binding for FOXA2, F0XM1, and ERα around introns, exons, and promoters. E) Known transcription factor binding motif enrichment near FoxM1, FoxA2, and ERα peaks in the presence of 17-p-estradiol. F) Known transcription factor binding motif enrichment near FoxM1, FoxA2, and ERα peaks in the presence of 17-β-estradiol compared to motif enrichment in the presence of vehicle only. N=5 for each condition.

Despite expectations that ERα would only interact with DNA upon E_2_ treatment, our data demonstrated that ∼84% of binding sites in E_2_-treated cells were also engaged in vehicle-treated cells (Figure 5A). Similarly, 46% of FOXM1 and 78% of FOXA2 binding sites in E_2_-treated cells were bound by ERα in vehicle-treated cells. In the presence of E_2_, an overlap of ∼46% between FOXM1 and ERα recruitment sites was observed, and ∼35% between FOXM1 and FOXA2. ∼55% of FOXM1:ERα co-bound sites were also bound by FOXA2. ∼25% of FOXM1 binding sites were bound by both FOXA2 and ERα. ∼71% of FOXA2 binding sites were also FOXM1 targets. ∼82% of ERα binding sites were also engaged by FOXM1 and ∼48% by FOXA2. These data suggest an overlap of binding between FOXM1, FOXA1, and ERα in β cells and an increase of co-binding in the presence of estradiol.

A large proportion of peaks for all three transcription factors was located in intergenic regions (Figure 5B-D), consistent with previous reports regarding ERα and FOXA2 binding-site locations (39,40), and in line with the substantial contribution of enhancers to β-cell gene regulation (41,42). We conducted known motif analysis of ChIP targets in both vehicle- and E_2_-treated cells. This evaluation revealed that all three transcription factors have enrichment for estrogen response element (ERE) and β-cell-enriched transcription factor binding motifs (Figure 5, E-F and data not shown). Despite the considerable overlap of binding sites between vehicle- and E_2_-treated cells, when using vehicle-treated ChIP binding sites as background, E_2_-treated cells still demonstrated enrichment of β-cell-specific transcription factor motifs, but not ERE, near FOXA2, FOXM1, and ERα binding sites (Figure 5F), suggesting that FOXA2, FOXM1, and ERα cooperate not only amongst themselves, but also with other transcription factors important for β-cell function.

## Discussion

We previously reported that FoxM1* protects against STZ- and cytokine-induced β-cell death. It also increases β-cell proliferation in aged male mice (16). In addition, FOXM1* increases β-cell Ca^2+^ influx in response to high glucose and improves glucose tolerance in young and aged males. In this study, scRNAseq demonstrated the upregulation of cell-cycle, apoptotic, and insulin-secretion genes in male β-FoxM1* β cells. Interestingly, *Glucose 6 phosphatase* (*G6pc2*) is among the genes involved in insulin secretion regulation upregulated in male β-FoxM1* β cells. G6PC6 negatively regulates basal insulin release (43) and could be at least partially responsible for normal fed and fasted glucose in β-FoxM1* males despite lowered glucose during a GTT, or may be elevated to counter-regulate other altered insulin secretion genes (16).

Expression of FoxM1* in human islets yielded increased insulin secretion only in male islets, prompting an investigation of the results of FOXM1* expression in female β cells. In contrast to males, female β-FoxM1* mice exhibited no increased β-cell proliferation or mass with age, nor did pregnancy induce elevated replication. In addition, FOXM1* did not improve glucose tolerance in female mice.

Because of the phenotypical similarities between mice lacking FOXM1 and ERα in the pancreas or β cell, and because FoxM1 and ERα interact in other tissues, we investigated whether increasing FOXM1 expression and activity in β cells could rescue glucose tolerance in ERα^Δβ^ mice. Like mice lacking ERα throughout the entire epithelium, ERα^Δβ^ mice exhibited glucose intolerance (9). However, in contrast to these mice, they also had elevated body weight, perhaps due to deletion in tissues outside the pancreas. They also exhibited elevated β-cell mass, contrary to our expectations given the reduced β-cell mass in ERα^-/-^ mice (12) in spite of their obesity (44). The increased β-cell mass in ERα^Δβ^ mice may have resulted from a failed attempt to respond to increased insulin resistance. In contrast, however, β-FoxM1*;ERα^Δβ^ mice exhibited a similar body weight as ERα^Δβ^ mice, yet had normal glucose tolerance and β-cell mass. This result suggests that increased FOXM1 activity can compensate for ERα.

The effects of estrogen on β cells are primarily mediated by ERα (35). Both nuclear and membrane-tethered ERα are present in β cells (11,45,46), yet the primary mechanism of ERα action investigated in the β-cell has been membrane signaling (9,46). Yet, several lines of evidence point toward the prevailing importance of non-nuclear ERα activity in the β cell. Membrane ERα signaling through extracellular signal-regulated kinase (ERK) induces Neurogenic Differentiation (NEUROD) binding and transactivation of *Insulin* (9). However, islets from female mice lacking either membrane or nuclear ERα functions show no defects in insulin secretion (47). These data, together with our data and previous reports demonstrating impaired glucose tolerance observed in female mice lacking both membrane and nuclear ERα in β cells, suggest that nuclear and membrane ERα can compensate for one another [Figure 4 and (9)].

Further supporting a role for nuclear functions of ERα, our ChIPseq experiments suggest some ERα activity occurs in an estrogen-independent or low-estrogen-responsive manner. Further corroborating this conclusion, male mice globally lacking membrane-tethered ERα but with nuclear ERα function have unaltered insulin secretion, while male mice globally lacking membrane-tethered ERα but with nuclear ERα function have unaltered insulin secretion (47). In addition, in most cell types, ERα has complimentary non-nuclear signaling and nuclear binding that coordinately regulate gene expression and other cellular functions.

This ChIP-seq was performed in βTC6 cells, and the sex of the mice used to make this cell line was not reported (48). However, in most tissues, including human β-cells, there is no difference in *Esr1* expression between the sexes, suggesting that differences in ERα activity are dependent on estrogen levels (49,50). This conclusion is also supported by phenotypical changes in male mice lacking ERα (47).

We investigated whether ERα and FOXM1 also bind the same sites in β-cell chromatin because of the functional overlap of ERα and FOXM1 in the β cell, and because their nuclear interactions in other cells are vital for driving the expression of particular gene sets (51). Our ChIPseq datasets revealed that, as in other cell types, FOXM1 and ERα displayed a high fraction of shared binding sites, and 17β-estradiol increased this proportion. Taken together, our data suggest that ERα binding facilitates FOXM1 interactions with chromatin to drive the transcription of genes involved in β-cell function, proliferation, and survival. Our data suggest a model whereby overexpression FOXM1* allows transactivation of β-cell gene promoters for which ERα facilitates FOXM1 binding. We propose a model where in control females, ERα binds near β-cell functional gene promoters and enhancers, allowing FOXM1 to bind (Figure 6A). However, in males and in females lacking ERα, wild-type FOXM1 is unable to access its targets (Figure 6B). However, the presence of an activated form of FOXM1 at levels higher than wild type overcomes the absence of ERα in males and in females lacking ERα (Figure 6C). This model is in agreement with other reports demonstrating that the presence of ERα can drastically alter the binding pattern of wild-type FOXM1 (51).

**Figure 6.**
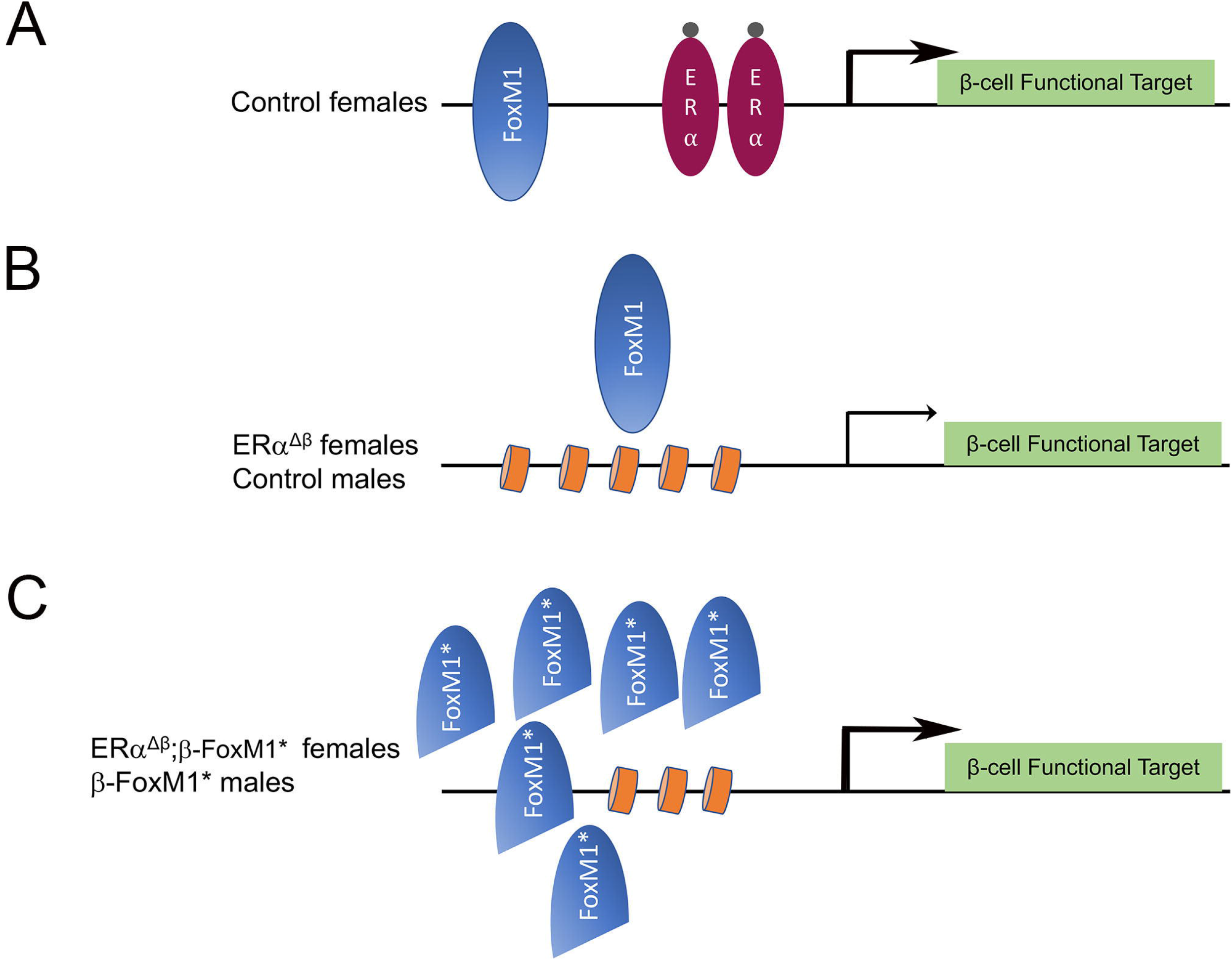
Working model for how activated FoxM1 improves β-cell function in mice lacking estrogen-dependent ERα activity in the β cell. A) In control females, ERα promotes binding of low levels of FoxM1 to β-cell functional target genes. B) Control males and female ERα mice lack ERα to aid in chromatin access for some target genes, preventing low levels of FOXM1 from binding. C) High concentrations of activated FOXM1 in β-FoxM 1* males and β-FoxM1*;ERα females allows FoxM1 to bind even in the absence of ERα.

Our results in female β-FoxM1 mice mirror the phenotype observed in mice lacking FOXM1 in the pancreas, where males exhibit impaired glucose tolerance or diabetes, but females have normal glucose tolerance except when further challenged metabolically (13,15). Our current data demonstrate that ERα binds DNA near many islet-enriched transcription factors, which is in line with previous reports of clusters of islet transcription factors (41). Also, ERα facilitates the genomic interactions of multiple other β-cell transcription factors important for β-cell function, which could explain why the loss of any single transcription factor that enhances insulin secretion, such as FOXM1, would have a negligible impact on β-cell function in females.

## Acknowledgments

We would like to thank Christopher Newgard and Thomas Becker (Duke Univeristy) for the gift of the FOXM1* adenovirus.

## Author Contributions

M.L.G. conceptualized the study, designed and performed experiments, and wrote and edited the manuscript. C.C. analyzed genomics data and wrote and edited the manuscript. G.P., E.M., K.K. and A.Y. designed and performed experiments.

